# Similar neural networks respond to coherence during comprehension and production of discourse

**DOI:** 10.1101/2021.06.24.449717

**Authors:** Matías Morales, Tanvi Patel, Andres Tamm, Martin J. Pickering, Paul Hoffman

## Abstract

When comprehending discourse, listeners engage default mode regions associated with integrative semantic processing to construct a situation model of its content. We investigated how similar networks are engaged when we produce, as well as comprehend, discourse. During fMRI, participants spoke about a series of specific topics and listened to discourse on other topics. We tested how activation was predicted by natural fluctuations in the global coherence of the discourse, i.e., the degree to which utterances conformed to the expected topic. The neural correlates of coherence were similar across speaking and listening, particularly in default mode regions. This network showed greater activation when less coherent speech was heard or produced, reflecting updating of mental representations when discourse did not conform to the expected topic. In contrast, regions that exert control over semantic activation showed task-specific effects, correlating negatively with coherence during listening but not during production. Participants who showed greater activation in left inferior prefrontal cortex also produced more coherent discourse, suggesting a specific role for this region in goal-directed regulation of speech content. Results suggest strong correspondence of discourse representations during speaking and listening. However, they indicate that the semantic control network plays different roles in comprehension and production.

## Introduction

When people comprehend discourse, they construct a mental model of its content (often termed a “situation model”) by integrating the incoming information with their prior semantic knowledge (Kintsch & van Dijk, 1978; Zwaan & Radvansky, 1998). These discourse comprehension processes engage regions in the default mode network (DMN), such as medial prefrontal cortex, precuneus, posterior cingulate, anterior temporal and lateral parietal cortex (Ferstl, Neumann, Bogler, & von Cramon, 2008; Yeshurun, Nguyen, & Hasson, 2021). This role in the construction of discourse models is supported by evidence implicating the DMN in semantic processing in general (Binder & Desai, 2011; Binder, Desai, Graves, & Conant, 2009) and in integration of multiple semantic cues in particular (Lanzoni et al., 2020). More broadly, the role of the DMN in the highest levels of language comprehension fits with data indicating that this network occupies the endpoint of a functional gradient in the brain that varies from low-level sensorimotor processes to high-level multimodal abstract thought (Margulies et al., 2016).

The DMN’s response to variation in the coherence of language input is a major source of evidence for its role in discourse-level processes. Coherence refers to the degree to which successive utterances relate meaningfully to each other and to the overall topic of the discourse (Gloser & Deser, 1992). Neuroimaging studies have typically manipulated discourse coherence by taking narratives and scrambling the order of the sentences so that they no longer tell a coherent story, or by constructing sets of sentences that are each individually meaningful but have no meaningful relationships between them. When presented with incoherent materials like this, it is impossible to construct a coherent situation model of the text. Accordingly, DMN regions show reduced activation to incoherent relative to coherent texts (Ferstl et al., 2008; Ferstl & von Cramon, 2002; Yarkoni, Speer, & Zacks, 2008) and DMN activation patterns become less strongly correlated across participants when narratives are incoherent (Lerner, Honey, Katkov, & Hasson, 2014; Lerner, Honey, Silbert, & Hasson, 2011; Simony et al., 2016).

Experimental manipulations of coherence have been important in establishing a role for the DMN in models of discourse content. These manipulations are, however, not representative of natural language. Healthy individuals never produce strings of entirely unrelated sentences when speaking. Instead, low coherence in natural language manifests as a weakening of the connections between utterances and an increased frequency of off-topic or tangential statements (Gloser & Deser, 1992; McNamara, Kintsch, Songer, & Kintsch, 1996). When listeners comprehend low coherence discourse of this type, they attempt to construct a situation model of its content but the process is made difficult by the fact that the model must be frequently updated and adjusted to accommodate shifts in topic and violations of the comprehender’s expectations (Kurby & Zacks, 2012; Zacks, Speer, Swallow, Braver, & Reynolds, 2007). Little is known about how the brain responds to natural discourse that is low in coherence but is not entirely incoherent. Some evidence from experimental manipulations of coherence suggests that activation in some parts of the DMN is *higher* for weakly coherent passages, relative to either highly coherent samples or completely incoherent material (Kuperberg, Lakshmanan, Caplan, & Holcomb, 2006; Maguire, Frith, & Morris, 1999). This is consistent with the idea that weakly coherent discourse places high demands on DMN regions that support the construction of mental models. One of the aims of the present study was to test whether a similar effect occurs in response to natural fluctuations in coherence in real samples of spoken discourse.

Situation models are thought to govern production as well as comprehension of discourse (Garrod & Pickering, 2004; Kintsch & van Dijk, 1978), but little is known about how these processes overlap in the brain. Few neuroimaging studies have investigated discourse production, but those that have reveal that generating discourse engages an extensive left-lateralized network including anterior temporal and inferior parietal DMN regions, as well as prefrontal regions that have been associated with planning and cognitive control (AbdulSabur et al., 2014; Awad, Warren, Scott, Turkheimer, & Wise, 2007; Blank, Scott, Murphy, Warburton, & Wise, 2002; Stephens, Silbert, & Hasson, 2010). Importantly, people activate a similar cortical network during the comprehension of discourse, suggesting a common neural basis (AbdulSabur et al., 2014; Awad et al., 2007; Silbert, Honey, Simony, Poeppel, & Hasson, 2014; Stephens et al., 2010; Wise & Geranmayeh, 2016). In particular, Pickering and Garrod (2004) have argued that, during conversation, interlocutors’ situation models (together with other linguistic representations) become aligned with each other (i.e., more similar), thus facilitating communication (Garrod & Pickering, 2004; Pickering & Garrod, 2021). Recent neuroimaging studies have provided evidence for this assertion. Studies measuring inter-subject correlations in neural activity indicate that speakers and listeners show correlated activation patterns in lateral parietal, and medial and lateral prefrontal cortex when speech is coherent (Heidlmayr, Weber, Takashima, & Hagoort, 2020; Silbert et al., 2014). This suggests a coupling of neural activity between individuals that facilitates communication (Silbert et al., 2014). Within individuals, however, it is not known whether producing speech places similar demands on model construction as comprehending it. A second aim of the present study was therefore to assess the degree to which neural correlations with coherence during comprehension are similar to those that occur when participants produce their own discourse. If the DMN plays a similar role in generating models for production as well as comprehension, we would expect to see similar effects.

In addition to representational processes which may be shared with comprehension, production of discourse relies on executive processes that regulate its content (Barker, Young, & Robinson, 2017; Kintz, Fergadiotis, & Wright, 2016). Executive control is necessary to ensure that coherence is maintained during speech production, such that discourse remains focused on the topic at hand and avoids irrelevant statements (Arbuckle & Gold, 1993; Marini & Andreetta, 2016). Supporting this view, performance on cognitive control tasks predicts the coherence of speech in older adults (Gold, Andres, Arbuckle, & Schwartzman, 1988; Kintz et al., 2016; Wright, Koutsoftas, Capilouto, & Fergadiotis, 2014) and deficits in coherence and narrative organization are commonly found in the speech of patients with impaired executive functions (Ash et al., 2006; Coelho, 2002; Hoffman, Cogdell-Brooke, & Thompson, 2020; Marini, Zettin, & Galetto, 2014). Recent evidence has identified a role for a specific aspect of executive function, namely semantic control, in the maintenance of coherence (Hoffman, Loginova, & Russell, 2018a). Semantic control processes regulate access to knowledge so that currently relevant concepts are retrieved and selected for use (Hoffman, McClelland, & Lambon Ralph, 2018b; Jefferies, 2013; Lambon Ralph, Jefferies, Patterson, & Rogers, 2017). Hoffman et al. (2018b) found that more coherent speech in young and older adults was predicted by better performance in a semantic control task that required participants to inhibit irrelevant semantic knowledge. In line with this finding, recent neuroimaging evidence links the activation of semantic control regions with the production of highly coherent discourse. Hoffman (2019) asked older adults to produce discourse on different topics. He found that activation increases in inferior frontal regions, implicated in semantic control (Badre & Wagner, 2007; Jackson, 2021), predicted the production of more coherent speech. Thus, while many discourse processes appear to be shared across production and comprehension, there is evidence that executive control networks may play specific roles in regulating content during production.

The aim of the present study was to investigate the neural correlates of coherence in naturalistic discourse passages, directly contrasting effects in speech comprehension with production. Previous studies have manipulated coherence using experimenter-designed discourse stimuli (Ferstl & von Cramon, 2002; Kuperberg et al., 2006). Here, we instead used a parametric approach to capitalise on the fluctuations in coherence that occur in natural speech. Participants underwent fMRI while producing and listening to discourse about a range of specified topics. We used previously validated methods from computational linguistics to quantify the global coherence of each passage of speech, i.e. the degree to which it related to the overall discourse topic (Gloser & Deser, 1992; Kintsch & van Dijk, 1978). We then investigated how neural activation correlated with variations in coherence.

As speech at the discourse level relies on widely distributed neural networks, we took a network-level approach to analysis. Traditionally, cortical networks have been identified in a discrete fashion, e.g., by using resting-state connectivity patterns to segregate the cortex into a set of distinct networks (e.g., Yeo et al., 2011). This approach assumes hard boundaries between networks. However, other theories propose that cortical function varies in a more continuous fashion, postulating graded variation in connectivity and function as one moves across the cortical surface (Lambon Ralph et al., 2017; Margulies et al., 2016). On this view, network membership is a graded property. In the present study, we used two approaches, one continuous and one discrete, to independently identify the DMN and other functional networks and to seek converging evidence for their roles.

In the first approach, we investigated effects of coherence along the continuous gradient of cortical organisation described by Margulies et al. (2016). Margulies et al. mapped the organisation of the cortex along a single dimension, such that regions of the brain that shared similar patterns of functional connectivity were located at similar points on the spectrum. At one end of this spectrum lie sensorimotor cortices, which are strongly functionally connected with one another. At the other end lie DMN-associated regions, which also show correlated patterns of activity but are anti-correlated with sensorimotor systems (see Figure 3B). Margulies et al. argued that the principal gradient reflects the organisation of cognitive processes in the brain, varying from stimulus-driven sensorimotor processes at one extreme to multimodal, internally generated thought at the other. We investigated the degree to which the brain’s response to coherence in discourse aligned with this gradient of neural organisation. In the second approach, we analysed the effect of coherence in three discrete networks of interest: the DMN, the semantic control network (SCN) and a network implicated in domain-general executive control (the multiple demand network or MDN; Fedorenko, Duncan, & Kanwisher, 2013). We tested how each of these networks responded to variations in the coherence of discourse and, in particular, the degree to which these effects were consistent for speaking and listening.

## Methods

### Participants

25 native English speakers from the University of Edinburgh participated in the study in exchange for payment. Their mean age was 24 (SD = 4.4, range = 18-35 years old) and they were all right-handed based on the Edinburgh Handedness Inventory (Oldfield, 1971). The study was approved by the Psychology Research Ethics Committee of the University of Edinburgh and all participants gave informed consent.

### Materials

In the production task, discourse was elicited by 12 prompts that asked about common semantic knowledge on particular topics (e.g., *Describe how you would make a cup of tea or coffee*; see Supplementary Materials for a complete list of prompts used in the production and comprehension tasks). Discourse production was contrasted with a baseline of automatic speech that involved the reciting of the well-known English nursery rhyme, Humpty Dumpty. For the comprehension task, we selected 24 samples of speech discussing 12 different topics. These were selected from a corpus of responses provided by participants in a previous behavioural study (Hoffman et al., 2018b). Speech comprehension used different topics to production, to avoid priming participants’ production responses with information presented in the comprehension trials. For each topic, we selected one highly coherent and one less coherent response for use in the study, to ensure sufficient variance in coherence across samples (highly coherent: M = 0.61, SD = 0.08; less coherent: M = 0.31, SD = 0.06; global coherence values, described in detail below). In addition, highly coherent and less coherent passages were of similar word length (highly coherent: M = 146, SD = 23.62; less coherent: M = 155, SD = 23.03; t = 1.33, *p* = 0.21). All comprehension speech passages were recorded by the same male native English speaker and their duration was 50 s each. For an example passage, see Supplementary Materials. The Humpty Dumpty rhyme was also pre-recorded as a comprehension baseline lasting 10 s.

### Design and procedure

Each participant performed two production and two comprehension runs, presented in an alternating sequence. Order of presentation was counterbalanced over participants, i.e., half of them began the experiment with a production run, whereas the other half began with a comprehension run. Each run was approximately 8 min and included six topics and five baseline trials, the order of which was fully randomised within each run. For both tasks, we created two sets of stimuli whose topics were matched for difficulty based on ratings obtained with a normative pre-test. In the pre-test, participants (N = 15), who were not part of the fMRI study, rated how difficult it would be to talk about each topic from 1 (easy) to 7 (difficult). The final sets of topics did not vary in difficulty rating (Production: M = 2.8, SD = 0.5; Comprehension: M = 2.4, SD = 0.76; *t*= 1.65, *p* = 0.11). In the comprehension task, half of the discourse passages were high coherent, while the other half were low coherent, counterbalanced over participants. This manipulation ensured that each participant was exposed to passages with a similar range of coherence scores, although we modelled coherence as a continuous parameter in our analyses.

The procedure for a single trial is shown in Figure 1A. Production trials started with the presentation of a written prompt on screen for 6 s. Participants were asked to prepare to speak during this period and to start speaking when a green circle replaced the prompt in the centre of screen. They were instructed to speak about the topic for 50 s, after which a red cross would replace the green circle. At this point participants were instructed to wait for the next prompt to appear on screen. The procedure for speech comprehension was the same except that participants were asked to listen to the speech attentively for 50 s while the green circle was on screen. For the baseline conditions, participants were instructed to recite (production task) or to listen (comprehension task) to the Humpty Dumpty rhyme for a 10 s period (in production they were asked to start again from the beginning if they reached the end of the nursery rhyme before the 10s had elapsed). The baseline conditions therefore involved grammatically well-formed continuous speech, but without the requirement to generate or understand novel utterances. The baseline trials were made shorter than the discourse trials to reduce boredom and to maximise the number of discourse trials in the study. All trials were presented with an interstimulus interval jittered between 3 s and 7 s (M = 5 s).

**Figure 1.**
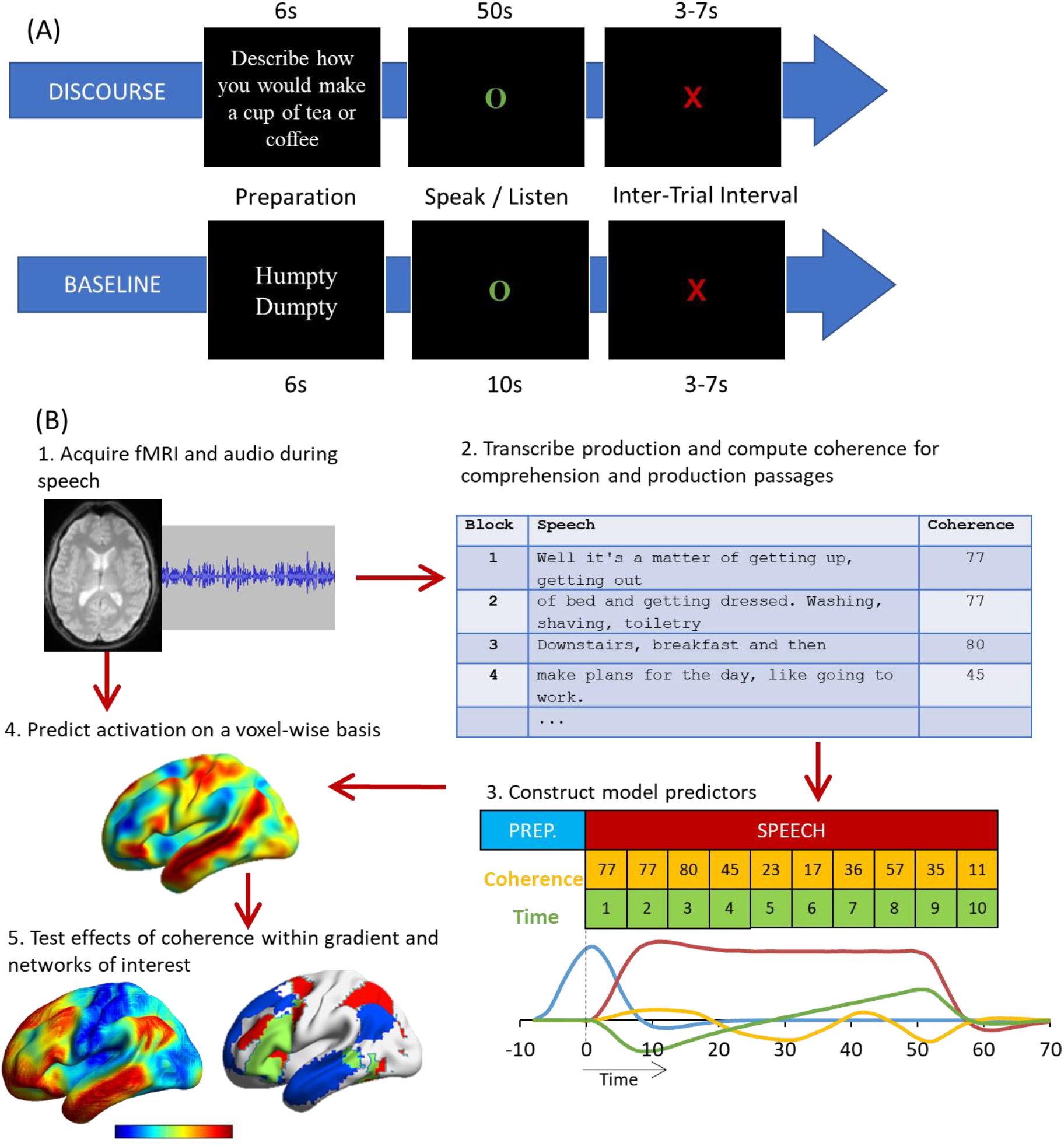
**A** – Illustration of a single trial of the discourse and baseline tasks. **B** – Stages in analysis.

Before scanning, participants were presented with two training trials to familiarise them with both tasks. They were also informed that they would receive a memory test after scanning to increase their motivation and attention during comprehension runs. In this test, participants answered 12 multiple choice questions, one for each topic presented during speech comprehension (each question had three choices). On average participants responded to 10/12 questions correctly (SD = 1.6; one-sample t-test comparing with chance performance: t = 31.21, *p* < 0.001). All participants responded above the chance level of 4/12. They were also asked to rate how well they could hear the speech samples in the scanner from 1 (inaudible) to 7 (perfectly audible), and rated auditability with a mean of 5.5/7 (SD = 1.0).

### Processing of speech samples

The stages in analysis are illustrated in Figure 1B. Responses to each prompt were digitally recorded with an MRI-compatible microphone and processed with noise cancellation software (Cusack, Cumming, Bor, Norris, & Lyzenga, 2005) to reduce noise from the scanner. They were then transcribed and the global coherence for each response was computed using computational analyses based on Latent Semantic Analysis (LSA) (Landauer & Dumais, 1997). In LSA, each word is represented as a high-dimensional vector in which words with similar meanings have similar vectors. Critically, the vectors for individual words can be combined linearly to represent the meaning of speech or text (Foltz, Kintsch, & Landauer, 1998). We used these representations of participants’ speech to quantify global coherence, the degree to which the utterances produced by a participant are related to topic they were asked about (Gloser & Deser, 1992). We measured this by calculating the similarity between utterances produced in response to a prompt and the prototypical semantic content expected for that prompt, in the following manner.

Global coherence was measured using the method first described in (Hoffman et al., 2018b). For each production response (target response), we began by computing an LSA representation for each participant’s response to the target prompt. Then these representations were averaged (excluding the target response) to obtain a composite vector for the prompt. This composite vector is therefore a semantic representation of a prototypical response to the target prompt, obtained by combining the responses of the whole group. Next, we divided the target response into a series of moving windows of 20 words in length, which each contained the current word and the 19 words preceding it. We computed an LSA vector for each of these windows. Then, these vectors were each compared to the prototypical vector using a cosine similarity metric that outputs a value ranging from 0 to 1. This value indicates how similar the speech produced in each window to the typical semantic content of responses to the same prompt. A high coherence value suggests that speech was strongly semantically related to the prompt being tested, whereas a low coherence value indicates speech that was semantically less related to the topic being probed. This method provided us with a moving estimate of coherence at the point each word was produced. To aggregate these values and relate them to the neural signal, we divided each 50 s speech period into blocks of 5 s and computed the mean global coherence for the words produced in each block (following the procedure in (Hoffman, 2019).

A similar approach was used to quantify global coherence for the speech samples presented in the comprehension task. Here, each speech window was compared with a composite prototype vector generated from the responses given to the prompt by 30 young adult participants in Hoffman et al. (2018).

### Image acquisition and processing

Participants were scanned on a 3T Siemens Prisma scanner using a 32-channel head coil. fMRI data were acquired at three echo times (13ms, 31ms, and 48ms) using a whole-brain multiband multi-echo acquisition protocol (Feinberg et al., 2010; Moeller et al., 2010; Xu et al., 2013). Data from these three echo series were weighted and combined, and the resulting time-series denoised using independent components analysis (ICA). This approach improves the signal quality in regions that typically suffer from susceptibility artefacts (e.g., the ventral anterior temporal lobes, (Kundu et al., 2017). The TR was 1.7 s and images consisted of 46 slices with an 80 × 80 matrix and isotropic voxel size of 3mm. Multiband acceleration with a factor of 2 was used and the flip angle was 73°. Four runs of 281 volumes (477.7s) were acquired. A high-resolution T1-weighted structural image was also acquired for each participant using an MP-RAGE sequence with 1mm isotropic voxels, TR = 2.5 s, TE = 4.6 ms.

Images were pre-processed and analysed using SPM12 and the TE-Dependent Analysis Toolbox 0.0.7 (Tedana) (DuPre et al., 2019). Estimates of head motion were obtained using the first BOLD echo series. Slice-timing correction was carried out and images were then realigned using the previously obtained motion estimates. Tedana was used to combine the three echo series into a single-time series and to divide the data into components classified as either BOLD-signal or noise-related based on their patterns of signal decay over increasing TEs (Kundu et al., 2017). Components classified as noise were discarded. After that, images were unwarped with a B0 fieldmap to correct for irregularities in the scanner’s magnetic field. Finally, functional images were spatially normalised to MNI space using SPM’s DARTEL tool (Ashburner, 2007).

Images were smoothed with a kernel of 8mm FWHM. Data were treated with a high-pass filter with a cut-off of 128 s and the four experimental runs were analysed using a single general linear model. Four speech periods were modelled as different event types: discourse production, baseline production, discourse comprehension, and baseline comprehension. Discourse periods were modelled as a series of concatenated 5 s blocks. This allowed us to include parametric modulators that coded the characteristics of speech: coherence of the speech, as computed in the earlier description, and time within the 50 s speech period. Time was included as a modulator because it is correlated with coherence: the later stages of a speech sample tend to be less coherent than the earlier ones (Hoffman, 2019). Modulators were mean-centred for each run. Additional regressors modelled the preparation periods for discourse and baseline in each task (6s events). Covariates consisted of six motion parameters and their first-order derivatives.

### Analyses

We first analysed the whole-brain activation for speech production and speech comprehension using the contrast discourse minus baseline in each task. For these analyses we employed a voxel-height threshold of *p* < 0.005 for one-sample t-tests with correction for multiple comparisons at the cluster level (*p* < 0.05 corrected), using SPM’s random field theory.

To investigate the effect of global coherence on activation during production and comprehension, we correlated effects with the principal connectivity gradient described in Margulies et al. (2016). This gradient provides a macroscale framework of functional and spatial cortical organisation, characterising a spectrum of activation that ranges from unimodal/sensory to multimodal/association areas associated with the DMN. Based on this connectivity gradient, we conducted two analyses using similar methods to Wang et al. (2020). First, we examined the effect of coherence on speech processing at different points on the gradient at the voxel level. We extracted the coherence effect (beta values) for production and comprehension in each voxel and computed voxelwise correlations between these values and the position of each voxel on the Margulies gradient. These correlations were performed in the whole brain and also within a mask of regions associated with semantic processing (defined as regions associated with the term “semantic” in the Neurosynth database; (Yarkoni, Poldrack, Nichols, Van Essen, & Wager, 2011). Second, to determine the reliability of these relationships across participants, we used linear mixed models to test how coherence varied along the cortical gradient. To do this, we divided the voxels along the gradient into 10 equally-sized bins (from bin 1 at the unimodal end to bin 10 at the multimodal end) and extracted a mean coherence effect for each participant for the voxels contained in each bin. We then conducted linear mixed models for production and comprehension to examine whether there was a linear (or higher order) relationship between the coherence effect and the connectivity gradient.

The gradient analysis tested how neural responses to discourse varied along a single, continuous dimension of neural organisation with the DMN as one of its endpoints. However, we were also interested in the effects on coherence on regions associated with semantic control and general executive function that fall at midpoints along the cortical gradient. To isolate effects in these regions more precisely, we extracted the mean effects of coherence from the following networks of interest (see Figure 5A):

1. The DMN, identified from Yeo et al’s (2011) 7-network parcellation of resting-state fMRI in 1000 participants.
2. The semantic control network (SCN), defined as significant voxels in Jackson’s (2021) meta-analysis. This meta-analysis identified regions that consistently show effects of cognitive control demands in semantic processing tasks.
3. The multiple-demand network (MDN), defined as regions responding to cognitive demands in multiple domains in Fedorenko et al. (2013). The MDN encompasses prefrontal and parietal and other regions (Duncan, 2010) and frequently acts in functional opposition to the DMN: while the DMN relates to more internally-oriented processes like mind-wandering and remembering the past, the MDN is present in more goal-oriented tasks (Mineroff, Blank, Mahowald, & Fedorenko, 2018).

As expected, many of the voxels in the SCN were also part of either the MDN or DMN (principally in lateral prefrontal cortex). To ensure independence between the three networks, voxels that fell within the SCN were excluded from the masks defining the DMN and MDN. Thus, MDN and DMN results reflect the functions of the portions of these networks that are *not* specifically implicated in controlled semantic cognition. A small number of voxels were also shared between DMN and MDN; these were excluded from the DMN mask.

To test effects of coherence, the mean coherence beta values for comprehension and production were extracted for each network for each participant. For each participant, we extracted data only from the top 20% of voxels that showed the largest effect in the relevant discourse-minus-baseline contrast (Tong et al., 2016). In other words, we restricted our analysis to those voxels within the network that were most strongly engaged by the discourse processing tasks. Coherence effect sizes were entered into a 2 × 3 (task x network) ANOVA.

Finally, we conducted an exploratory analysis of individual differences to determine whether brain activity during speech production differed between highly coherent and less coherent speakers. Following Hoffman (2019), we calculated the mean coherence for each participant over all of their responses and included this value as a covariate in the group-level analysis of the BOLD response. This allowed us to explore whether participants who tended to speak coherently about the topics provided showed different patterns of activation to those who were less coherent.

## Results

### Characteristics of speech

In speech production, participants produced 128 words to each prompt on average (range 120-145) and the mean global coherence was 0.49 across all responses (range 0.43-0.66).

We divided each 50 s speech period into ten segments of 5 s each to examine the variation of global coherence over time in speech production and in speech presented for comprehension. For production, we observed a high negative correlation between global coherence and the position of segments in the 50s discourse period (r = -0.9) (see Supplementary Materials). In other words, the longer participants spoke for, the less related their speech became to the prescribed topic. Coherence and time segments were also strongly correlated in speech presented in comprehension runs (r = -0.93). To avoid a confounding effect of this factor, we included segment position as a covariate in the brain analysis for speech production and comprehension, as in Hoffman (2019) (see Figure 1B). We found that the number of words produced in each segment did not vary over time (r = 0), indicating that participants did not slow their speech rate or run out of things to say during these later periods of speech. These patterns were similar in speech samples used for comprehension. Finally, there was no correlation between number of words in a segment and its coherence, for either production (r = -0.2, *p* = 0.58) or comprehension (r = 0.3, *p* = 0.34).

### Activation during speech comprehension and speech production

Figure 2 shows activation for discourse production and comprehension against automatic speech baselines. For production, whole-brain analysis revealed a left-biased activation pattern that included inferior frontal gyrus (pars triangularis and pars opercularis), angular gyrus, frontal orbital cortex, the middle frontal gyrus, the anterior temporal lobe and the length of the superior temporal sulcus. A strikingly similar pattern was found in the comprehension task, but with more activation in left posterior temporal regions and additional activation in the right anterior temporal lobe. This is consistent with the semantic-related network found in other studies relating to discourse processing (e.g., (Hoffman, 2019; Silbert et al., 2014) and indicates a high degree of overlap in the networks recruited for high-level discourse comprehension and production.

**Figure 2.**
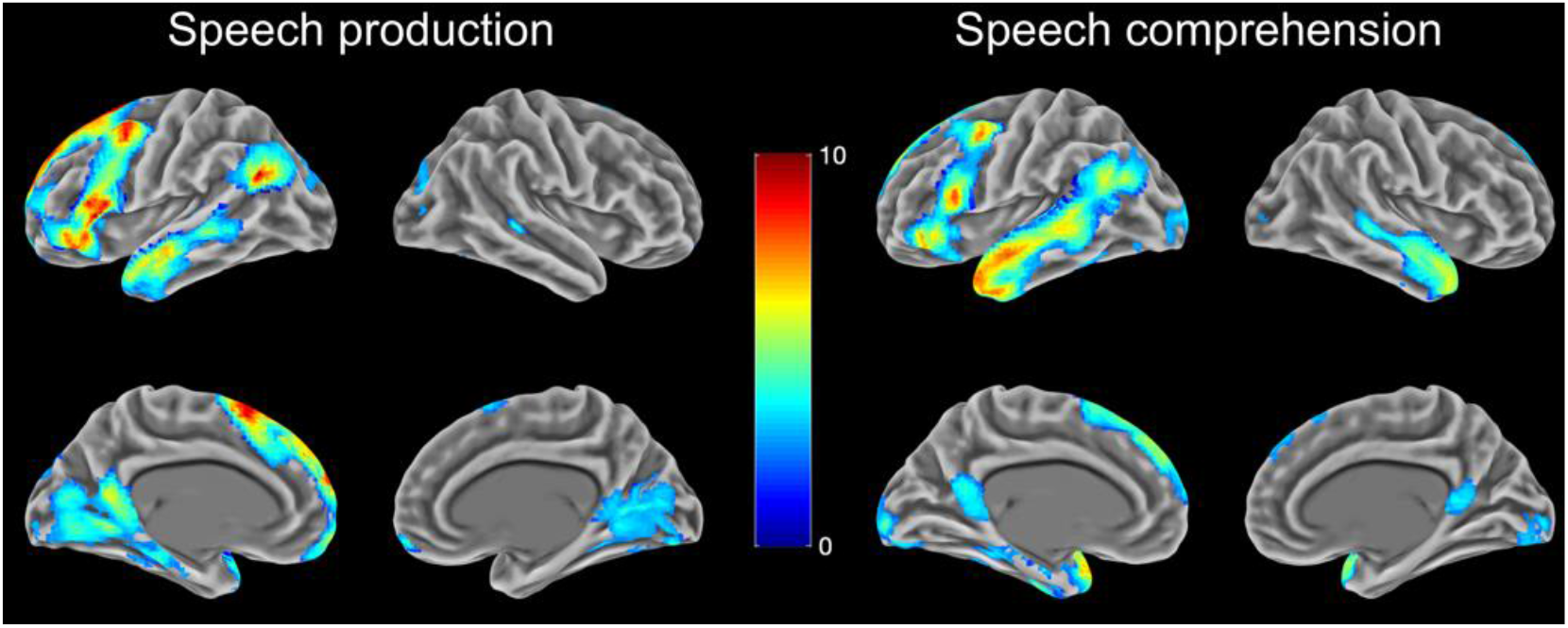
Neural activation during speech comprehension and speech production over automatic speech baseline. Images thresholded at cluster-corrected *p* < 0.05.

### Activation during speech comprehension and speech production as a function of coherence

Coherence effects across the whole brain are shown in Figure 3A. Activation in blue areas was positively correlated with coherence, increasing when participants produced or comprehended more coherent speech, whereas red areas were negatively correlated with coherence, showing greater activation when speech was less coherent. For speech production, positive effects of coherence were found principally within the supramarginal gyrus, posterior inferior and middle temporal gyri and the precentral gyrus. In contrast, negative effects of coherence were found predominantly in DMN regions such as the angular gyrus, anterior middle and superior temporal gyri, the posterior cingulate and ventromedial prefrontal cortex. A similar set of regions showed negative effects of coherence in speech comprehension, with a particularly strong effect in the lateral temporal cortices.

**Figure 3.**
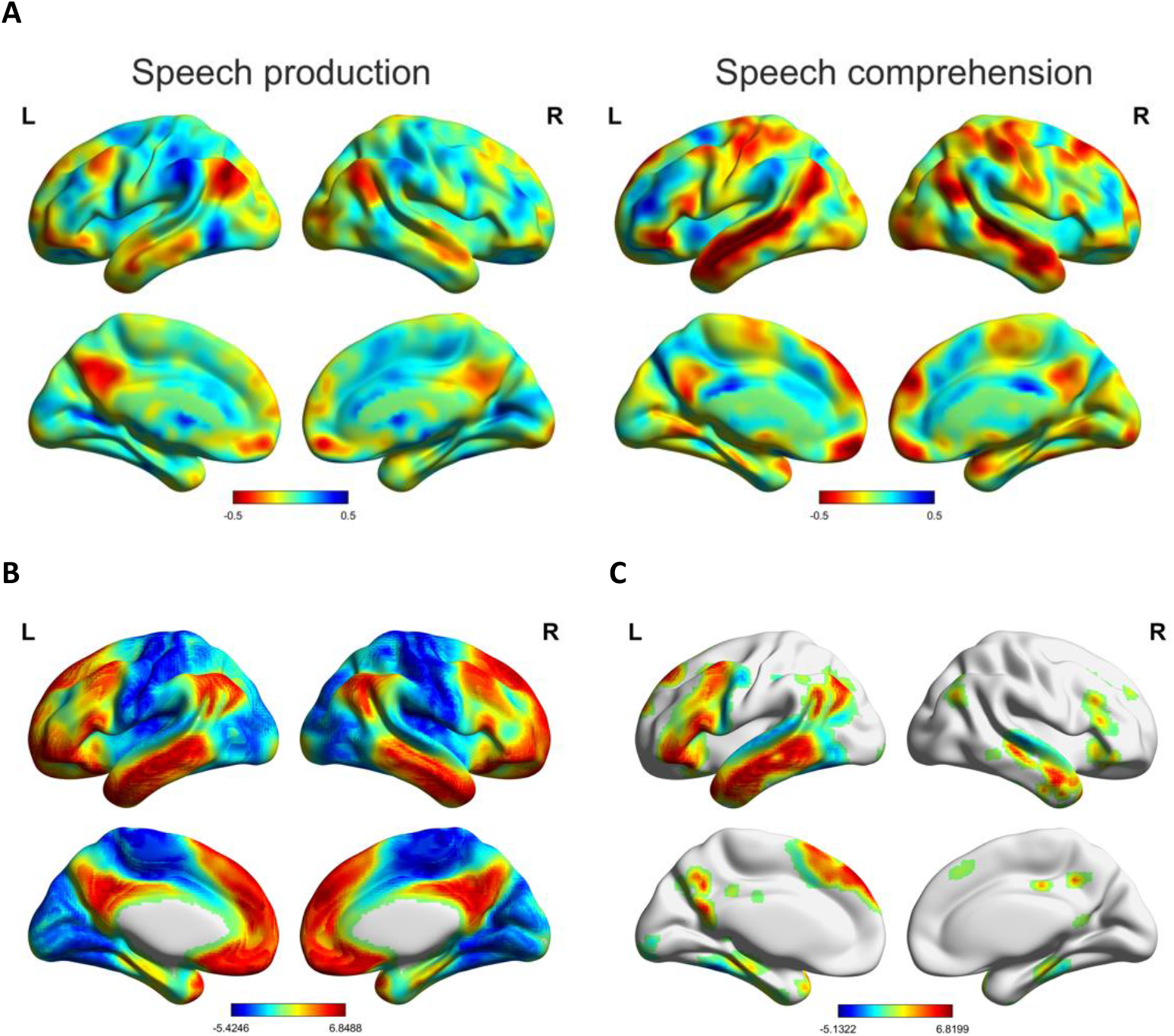
**A** - Unthresholded maps of the coherence effect (beta values) on speech production and speech comprehension. **B** - Principal connectivity gradient map from Margulies et al. (2016), with the unimodal extreme of the gradient shown in blue and the DMN extreme in red. **C** – Areas of the Marguiles et al. gradient falling within the mask of semantic processing regions.

Figure 3B shows the principal functional gradient reported by Marguiles et al. (2016), which arranges regions along a continuum from primary sensory and motor regions to the multimodal cortex of the DMN. Visual inspection suggests alignment between this gradient and the effects of coherence: areas towards the DMN end of the cortical gradient appear to show the strongest negative response to coherence. We assessed this pattern formally in the next section.

We analysed our data in two different ways. First, we correlated coherence effects with the connectivity gradient at the voxel level (Wang, Margulies, Smallwood, & Jefferies, 2020). Correlation tests were conducted between the cortical gradient values and group-mean beta values of the coherence effect in production and comprehension (Figure 4, panels A-B, D-E). For production, we found a negative correlation between the coherence effect and the principal gradient (r = -0.4, p < 0.001). The correlation was stronger when the analysis was restricted to semantic regions only (r = -0.5, p < 0.001; voxels shown in Figure 3C). The closer voxels were to the multimodal, DMN end of the gradient, the more likely they were to show a strong negative correlation with coherence. For comprehension, activation as a function of coherence was weakly correlated with the gradient at the whole-brain level (r = -0.09, p < 0.001), but more strongly correlated when analysing only voxels within semantic areas (r = - 0.35, p < 0.001). However, Figure 4D also suggests the possibility of a non-linear relationship between coherence and the gradient at the whole-brain level, in which voxels at both extremes of the gradient show greater activation for less coherent speech.

**Figure 4.**
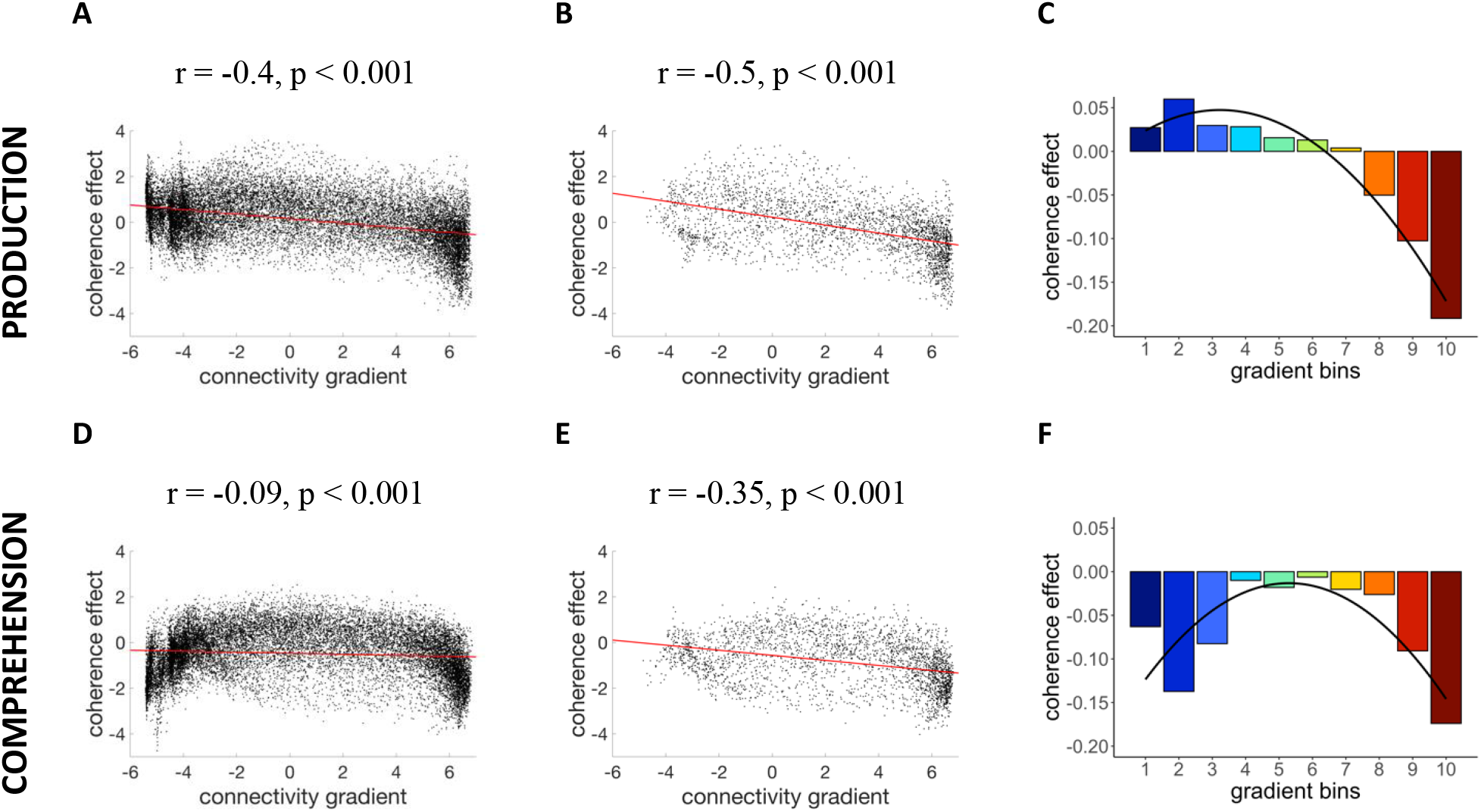
**A** and **B** – Correlations between coherence beta values and the connectivity gradient in production at the whole-brain level and within semantic regions, respectively. **C** – Model fit for the mixed effects model that examined the effect of the gradient on coherence activation during speech production. **D** and **E** – Correlations between the coherence beta values and the connectivity gradient in comprehension at the whole-brain level and within semantic regions. **F** – Model fit for the mixed effects model that tested the effect of the gradient on coherence activation during speech comprehension.

Next, to account for the possibility of a non-linear relationship between coherence and the gradient, we segmented it into ten bins from unimodal (bin 1) to multimodal DMN regions (bin 10) (Wang et al., 2020). Each voxel was assigned to one of the bins based on its position on the gradient and the coherence effects in the voxels for each bin were averaged for each participant. We then fitted linear mixed models predicting these coherence values for comprehension and production separately (Figure 4, panels C and F). Our fixed effect was the number of the gradient bin (from 1 to 10), treated as a numeric variable and centred. To test for higher order relationships, we included a quadratic term for gradient bin in all models as this predictor improved model fits (p < 0.05) over and above the linear term alone (see Supplementary Materials for model comparisons). All models included random intercepts and slopes by participant and p-values were obtained with the lmerTest package (Kuznetsova, Brockhoff, & Christensen, 2017) via Satterthwaite’s method.

We found a significant linear effect for speech production (β = -0.022, SE = 0.007, t = -2.946, p = 0.007). As shown in Figure 4C, voxels towards the sensorimotor end of the gradient showed greater activation when speech was more coherent, while those towards the DMN end showed more activation when participants produced less coherent speech. The quadratic term did not reach significance for the production task (β = -0.005, SE = 0.003, t = -1.789, p = 0.086). For speech comprehension, we found no linear relationship between the coherence effect and the gradient (β = -0.002, SE = 0.011, t = -0.220, p = 0.828) but the quadratic term showed a significant effect (β = -0.006, SE = 0.003, t = -2.116, p = 0.045). As shown in Figure 4F, there was a strong negative effect of coherence at the sensorimotor end of the gradient, which gradually decreased at middle parts of the spectrum before increasing at the DMN end. Thus, comprehension and production effects converged in DMN regions, where there was greater activation for less coherent discourse, but were less consistent in sensorimotor regions.

### Coherence effects in networks of interest

To confirm and extend the results found for the cortical gradient, we compared the effects of coherence across language tasks in regions belonging to the DMN, SCN and MDN (see Figure 5). We computed a 2 × 3 (task x network) ANOVA, revealing a main effect of network (*F* (2, 48) = 10.2, *p* < 0.001) and a task × network interaction (*F* (2, 48) = 3.62, *p* = 0.034). Post-hoc t-tests with Holm-Bonferroni correction indicated that, during comprehension, DMN and SCN exhibited similar effects of coherence (*p* = 0.38) which were more negative than those seen in the MDN (*p* < 0.05). In production, however, effects in DMN and SCN differed from one another (*p* = 0.04). MDN differed from DMN (*p* = 0.005) but not SCN (*p* = 0.64). Thus, the DMN showed activation increases during low coherence speech that were consistent across tasks, while less coherent discourse produced greater activation in SCN regions only in comprehension. There were no effects of coherence in MDN regions that were not specifically associated with control over semantic processing.

**Figure 5.**
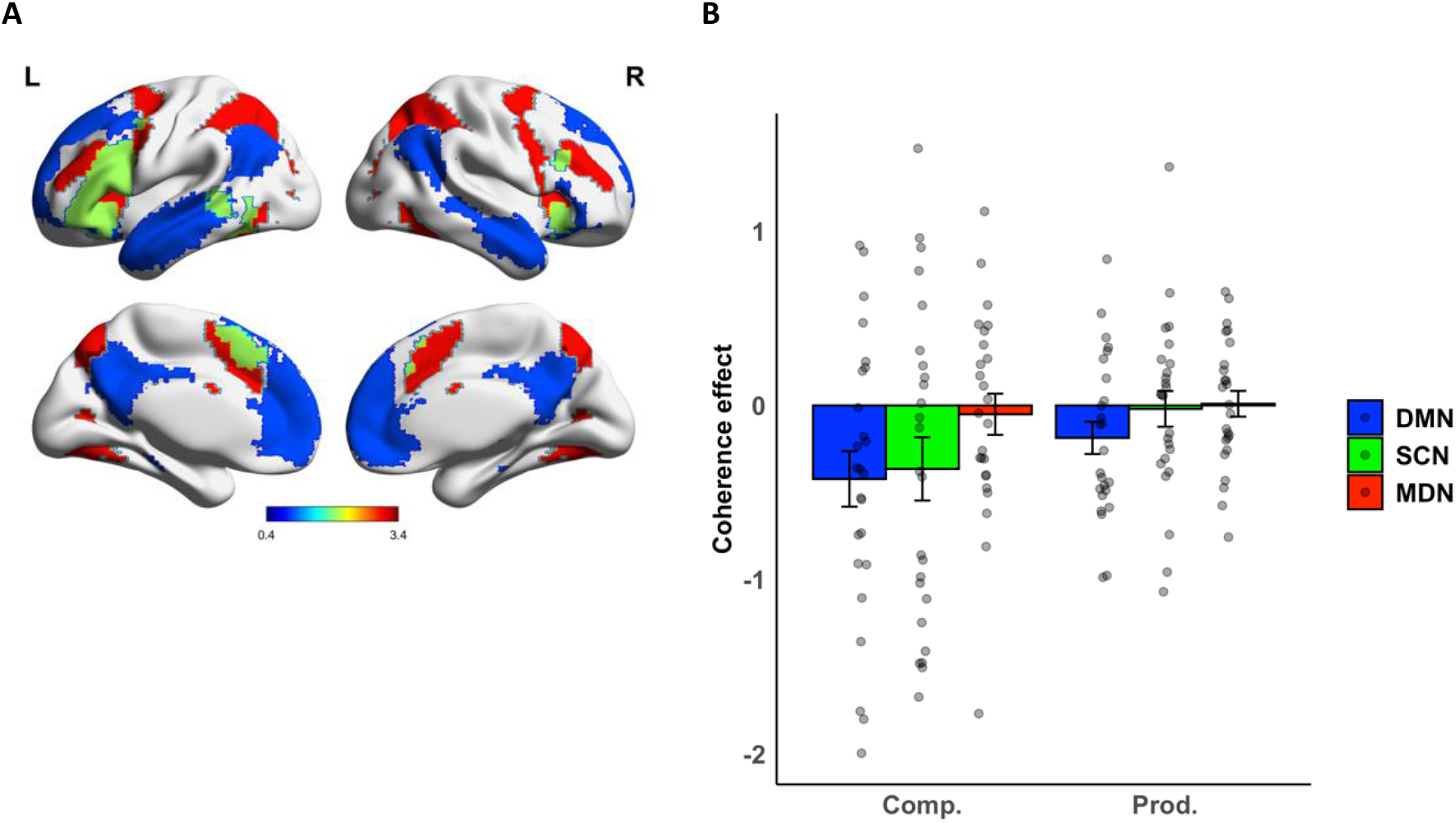
**A** – Masks used for the default mode network (DMN - blue), semantic control network (SCN - green), and multiple demand network (MDN - red). **B** – Effects of coherence in each network.

### Effects of individual differences in coherence on activation during speech production

The prior within-subject analyses all investigated how activation covaried with fluctuations in the coherence of speech over time. To explore how activation differed between more and less coherent speakers, we included the mean coherence of the speech produced by each participant as a predictor of activation. Activation during speech production was positively correlated with the coherence of the speaker in a cluster of voxels in the pars triangularis portion of the left inferior frontal gyrus (17 voxels at uncorrected *p* < 0.001 with a peak at [-54, 36, 12], *t* = 5.08). Participants who produced more coherent speech showed greater activation in this area while speaking. This effect did not survive correction for multiple comparisons across the whole brain; however, it is consistent with previously-found associations of this area with coherence (Alyahya, Lambon Ralph, Halai, & Hoffman, submitted for publication; Hoffman, 2019) and with the theory that the semantic selection processes served by the inferior frontal gyrus are important for maintaining coherence (Hoffman et al., 2018). No other regions showed a positive association with coherence at *p* < 0.001. Less coherent speakers showed more activation in the cerebellum (50 voxels at uncorrected *p* < 0.001 with a peak at [-3, -39,-15], *t* = 4.37).

## Discussion

In this fMRI study, we investigated how neural activity covaried with the coherence of discourse during speech production and comprehension. We used a global measure of coherence which tracked the degree to which speech conformed to the expected discourse topic (Gloser & Deser, 1992; Hoffman et al., 2018b). DMN regions showed activation increases when discourse was lower in coherence, whether participants were listening to another speaker or producing their own narratives. The brain’s response to coherence was closely aligned with a cortical connectivity gradient that arranges brain regions on a continuum from sensorimotor processing to multimodal abstract thought (Margulies et al., 2016). The multimodal end of this gradient, most associated with the DMN and with integrative higher-order cognition, showed the strongest increases for lower coherence in speech. Coherence effects in comprehension and production diverged in the semantic control network. Regions involved in semantic control showed negative coherence correlations in comprehension that did not occur during production. Our results have implications for understanding the shared processes that support the representation of discourse during speaking and listening, and the points at which these processes diverge.

When listening, participants showed greater DMN activation when discourse was less coherent. This result is consistent with the hypothesized role of this network in constructing situation models that represent the elements of discourse, their meanings and their relations with one another (Baldassano et al., 2017; Heidlmayr et al., 2020; Lerner et al., 2011). We begin by noting that, unlike many previous studies, we did not use speech passages which were entirely incoherent (e.g., sentences in a scrambled order). Under these conditions, DMN activation declines (Ferstl et al., 2008; Ferstl & von Cramon, 2002; Yarkoni et al., 2008), presumably because attempts to build a model of the discourse are abandoned. In contrast, we assume that when people listened to the speech in our study, they always attempted to form a situation model by integrating the incoming speech input with their prior knowledge and expectations of the topic under discussion. When the speech input was less coherent, we assume that this mental representation required frequent updating and reconfiguration, resulting in increased DMN activation. This interpretation is supported by evidence that DMN activity increases when participants encounter event boundaries in narratives and have to update their mental representations (Speer, Zacks, & Reynolds, 2007; Whitney et al., 2009). Our results are consistent with previous evidence showing that DMN areas are involved in constructing coherent mental models of discourse (Ferstl et al., 2008; Ferstl & von Cramon, 2002; Heidlmayr et al., 2020; Jacoby & Fedorenko, 2020; Kuperberg et al., 2006; Yeshurun et al., 2021). Uniquely, however, our study reveals that similar effects are present when people produce their own discourse. This suggests that the people experience similar difficulty in developing less coherent situation models to use as the basis for their own utterances as when they are comprehending other people’s utterances.

Overall, we found that listening and speaking both activated a left-dominant network of frontal, temporal, and inferior parietal regions, consistent with shared activation patterns across comprehension and production in previous studies (e.g., e.g., Silbert et al., 2014). Beyond mere coactivation, however, the present study reveals that much of this network responds to the content of speech in similar ways across comprehension and production. Specifically, participants activated DMN regions more when their own discourse was less related to the topic of discussion. This negative effect could partly be a consequence of participants’ self-monitoring of their own utterances. If participants monitored and updated their situation models on the basis of their speech output, this would result in greater DMN activation during passages of less coherent speech production, just as it does during comprehension. It is likely, however, that situation models also play a much earlier role in planning and generating speech content. Studies of discourse comprehension suggest that increases in DMN activation are indicative of major shifts in the content of situation models (Speer et al., 2007; Whitney et al., 2009). When such shifts occur during production, the likely consequence is speech that lacks coherence and fails to conform to the prescribed topic.

Overall, then, our results suggest that the mechanisms involved in constructing situation models are largely shared between production and comprehension. Previous studies have provided support for this idea by comparing the brain activation of the person producing an utterance with that of the comprehender (Heidlmayr et al., 2020; Silbert et al., 2014; Stephens et al., 2010). In contrast, here we have shown that convergence occurs within individuals when comparing their activation profile as a speaker and as a listener. Our results can be explained by models of language processing which assume that semantic representations are shared between comprehension and production (Gambi & Pickering, 2017; Hagoort, 2013; Kintsch & van Dijk, 1978; Levelt, Roelofs, & Meyer, 1999). This shared representation may allow the production system to make forward predictions that aid comprehension (Dell & Chang, 2014; Pickering & Garrod, 2007) and help interlocutors to align their situation models during conversation (Garrod & Pickering, 2004).

Our interpretation of our results has focused on the idea that the DMN supports situation models of discourse content. This notion is highly compatible with other theories implicating this network in representing event characteristics during episodic recall (Ranganath & Ritchey, 2012) and with the notion that inferior parietal cortex in particular functions as a multimodal buffer of recent experience (Humphreys & Lambon Ralph, 2015). These theories share the view that the DMN generates mental models of events and situations, whether these draw from current experience, recall from memory or from processing discourse. Thus the role of the DMN is not limited to verbal processing; indeed, DMN responses are influenced by manipulations of coherence when people watch movies as well as when they listen to spoken stories (Lerner et al., 2011).

Another set of researchers have investigated the involvement of DMN in mind-wandering and internally generated thoughts (Andrews-Hanna, Smallwood, & Spreng, 2014; Christoff, Gordon, Smallwood, Smith, & Schooler, 2009). These studies are consistent with the view that the DMN represents mental models of events and situations, as these are a common constituent of self-generated thought (Andrews-Hanna et al., 2014; Binder et al., 1999). Increases in DMN activity have also been associated with task-unrelated thoughts and attention lapses, potentially reflecting unwanted engagement of self-generated thoughts (Christoff et al., 2009; Mason et al., 2007). In our study, the DMN appeared to be making an active contribution to task performance as large parts of the network were *positively* engaged by the discourse tasks (see Figure 2). Is it possible, nevertheless, that the effects of coherence we have observed reflect the intrusion of self-generated thoughts that are not relevant to the task at hand? During comprehension, this account would imply that participants experienced more task-unrelated thoughts during less coherent passages of speech. Although we cannot rule out this possibility, we consider it unlikely since making sense of less coherent speech requires greater attentional focus, not less. During the production task, participants generated their own speech and here it is likely that irrelevant thoughts would lead to the production of a weakly coherent response.

Emerging evidence suggests that cognitive control networks are critical for task-based regulation of DMN activity (Andrews-Hanna et al., 2014). With respect to discourse production in particular, previous studies have found that participants with better executive and semantic control abilities produce more coherent speech (Hoffman et al., 2018b; Wright et al., 2014). In the present study, evidence for this executive regulation is mixed. In an analysis of individual differences, we found that the more coherent speakers in our study activated a left inferior prefrontal region (BA45) more than those who were less coherent. This region is the dominant node in a network of semantic control regions and is specifically implicated in inhibition of irrelevant semantic knowledge (Badre & Wagner, 2007; Jefferies, 2013). This finding supports the idea that semantic control regions regulate the selection of ideas during speech production. When we looked at neural correlations within participants, we found the SCN’s association with coherence was task-dependent. During comprehension, less coherent speech activated the SCN more, in line with the idea that less predictable speech input requires greater executive control. In production, this negative effect was not present. However, we did not find the positive association with coherence that the executive regulation account would predict, and which we found in our previous study in older adults (Hoffman, 2019). Thus, there is mixed evidence as to whether greater SCN engagement leads to the production of more coherent discourse and more research is needed to investigate how and when such regulation occurs.

Beyond the SCN, we found no effects of discourse coherence in the MDN that supports domain-general cognitive control. Some researchers have proposed a role for domain-general MDN regions in regulating speech production (Wise & Geranmayeh, 2016). Here, however, we found these regions were not modulated by the coherence of speech when either speaking or listening. These results suggest that involvement in discourse processing is restricted to those parts of the MDN that are specifically implicated in control over semantic processing (which, in our analysis, were assigned to the SCN instead). This aligns with the general view that cognitive control at the higher levels of language processing draws on specific neural resources distinct from those engaged in other cognitive domains (Fedorenko, Behr, & Kanwisher, 2011; Mineroff et al., 2018).

Our results add to an emerging body of literature examining how effects of semantic processing vary along a cortical gradient from sensorimotor regions to the DMN. Other studies have found that when processing word pairs, pairs with a stronger semantic relationship cause greater activation in areas towards the DMN (Gao et al., 2021; Wang et al., 2020). There are many differences between word-pair semantic judgements and the more natural form of comprehension studied here. However, one potential way of integrating these findings is to assume that semantically related word pairs bring to mind a mental model of particular situation or context, which is supported by the DMN.

One major contribution of the present study is to extend findings relating to comprehension into the domain of discourse production. Although we found convergence in cortical regions towards the DMN end of the cortical gradient, coherence effects in speaking and listening appeared to dissociate at the sensorimotor extreme of the principal cortical gradient. These regions showed more activation to less coherent discourse in comprehension but not in production. We suggest that these effects may relate to lower levels of speech perception. It is well-known that when perceiving speech in noisy conditions, such as those experienced in a MRI scanner, processing is easier when speech content is highly semantically coherent (Boothroyd & Nittrouer, 1988). Accordingly, activation in auditory cortices is increased when people hear less semantically predictable words during narrative comprehension (see also see also Friederici, Rüschemeyer, Hahne, & Fiebach, 2003; Willems, Frank, Nijhof, Hagoort, & van den Bosch, 2016). Thus, the enhanced response of sensorimotor regions to less coherent passages may reflect the greater demands that these place on speech perception systems.

In conclusion, we found increased DMN activation when participants produced and listened to less coherent discourse. This result suggests that people update and reconfigure representations of discourse content in similar ways during production and comprehension. However, our results also revealed activation increases in semantic control regions during the comprehension of less coherent discourse, which were not present during production. This indicates that the relationship between coherence and executive control systems is task-dependent. Thus, our findings contribute to understanding of the ways neural processes are shared between the production and comprehension of discourse, but also identify the points at which they diverge.

## Data and code availability

Analysis code and study data are available at: https://osf.io/nwbk3/

## Acknowledgements

PH was supported by a BBSRC grant (BB/T004444/1). We are grateful to Beth Jefferies and Xiuyi Wang for helpful discussions regarding these data. Imaging was carried out at the Edinburgh Imaging Facility (www.ed.ac.uk/edinburgh-imaging), University of Edinburgh, which is part of the SINAPSE collaboration (www.sinapse.ac.uk). We are grateful to the University of Minnesota Center for Magnetic Resonance Research for sharing their neuroimaging sequences.

## Notes

https://osf.io/nwbk3/

### Competing Interest Statement

The authors have declared no competing interest.

