## Supplementary Materials for "Similar neural networks respond to coherence during comprehension and production of discourse" (from another version of this preprint: version 1, 2021-06-24)

**1. List of prompts used in production and comprehension tasks.**

**1.1. Production.**

1. Describe how you would make a cup of tea or coffee.

2. Why are some people concerned about climate change?

3. Why is it important to have a balanced diet?

4. Describe the steps you’d take to order-in food.

5. Describe a typical visit to a grocery store.

6. How would you prepare to go on holiday?

7. Do you think the internet has improved people's lives?

8. What would you recommend doing during a job interview?

9. What do people usually do on New Year’s Eve in the UK?

10. What sorts of things do people do to cope with stress?

11. What sort of things does a teacher do when at work?

12. What sort of things usually happen at a wedding?

**1.2. Comprehension.**

1. What would it be like to live in Antarctica?

2. What happens when a storm is forecast in the UK?

3. What do the police do when a crime has been committed?

4. Which is your favourite season and why?

5. What do you like or dislike about Christmas?

6. Do you think it’s a good idea to send people to live on Mars?

7. Why do people come to Scotland on holiday?

8. What sort of things do you have to do to look after a dog?

9. Describe a typical visit to a restaurant.

10. What are the advantages and disadvantages of going to university?

11. What do people usually do when getting ready for work in the morning?

12. Describe the steps you would need to take if going somewhere by train.

**2. Example of passage used in the comprehension task.**

**Prompt: What would it be like to live in Antarctica?**

“I don’t know whether Antarctica is north or south but I imagine that it’s cold. It’s one or the other. I don’t know if there are penguins there. But there would be penguins around, I guess. If there were penguins there they would in the Antarctic with you. You’d probably be living in an igloo, riding reindeers and stuff and killing seals to survive and wearing polar bear hide or something weird. I don’t know. You’d probably be pretty chilly all the time, so you’d have to layer up a lot. Beware of killer whales that burst through the ice and eat you in your sleep. You’ve got to really just be aware of the dangers of nature. Also snowstorms. They can pretty killer. If you get stuck in a snowstorm you’re pretty screwed because they can be pretty harsh”

**3. Correlations between global coherence and temporal segment.**

| **Production** | **Comprehension** |
| --- | --- |
| 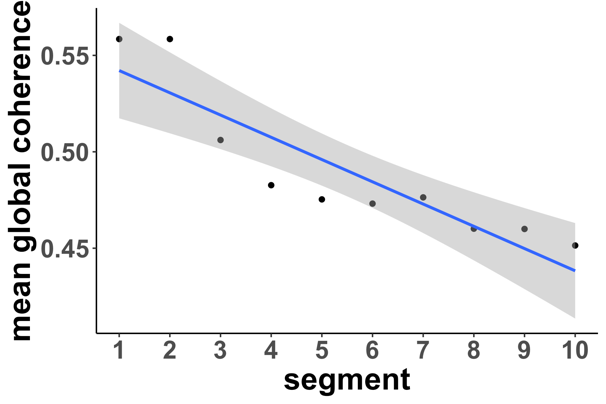 | 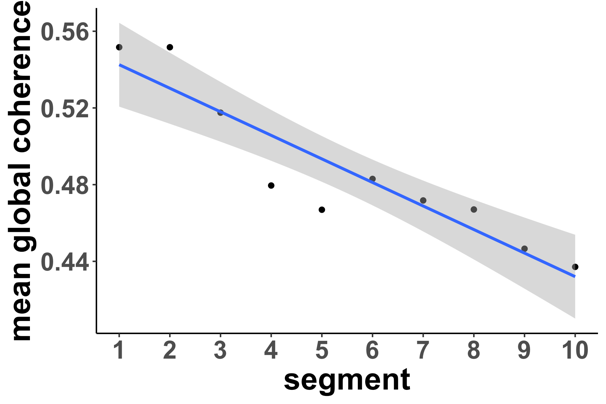 |

**4. Model comparisons for linear mixed models in production and comprehension tasks.**

**4.1. Production**

|  | **Df** | **AIC** | **BIC** | **logLik** | **deviance** | **Chisq** | **Chi Df** | **Pr(>Chisq)** |
| --- | --- | --- | --- | --- | --- | --- | --- | --- |
| coherence effect ~ (1\|subj) | 3 | -10.57 | -0.01 | 8.29 | -16.57 |  |  |  |
| coherence effect ~ linear bin + (1\|subj) | 4 | -35.25 | -21.16 | 21.62 | -43.25 | 26.68 | 1 | 0.00 |
| coherence effect ~ linear bin + quadratic bin + (1\|subj) | 5 | -42.21 | -24.61 | 26.11 | -52.21 | 8.96 | 1 | 0.00 |
| coherence effect ~ linear bin + quadratic bin + cubic bin +(1\| subj) | 6 | -41.19 | -20.06 | 26.60 | -53.19 | 0.98 | 1 | 0.32 |

**4.2. Comprehension**

|  | **Df** | **AIC** | **BIC** | **logLik** | **deviance** | **Chisq** | **Chi Df** | **Pr(>Chisq)** |
| --- | --- | --- | --- | --- | --- | --- | --- | --- |
| coherence effect ~ (1\|subj) | 3 | 100.92 | 111.48 | -47.46 | 94.92 |  |  |  |
| coherence effect ~ linear bin + (1\|subj) | 4 | 102.70 | 116.78 | -47.35 | 94.70 | 0.22 | 1 | 0.64 |
| coherence effect ~ linear bin + quadratic bin + (1\|subj) | 5 | 96.28 | 113.88 | -43.14 | 86.28 | 8.42 | 1 | 0.00 |
| coherence effect ~ linear bin + quadratic bin + cubic bin +(1\| subj) | 6 | 95.59 | 116.72 | -41.80 | 83.59 | 2.68 | 1 | 0.10 |

**5. Summary tables of linear mixed models in production and comprehension tasks.**

**5.1. Fixed effects in production.**

| **LMM for speech production: coherence effect ~ linear bin + quadratic bin + (1\|subj)** | | | | | |
| --- | --- | --- | --- | --- | --- |
|  | **Estimate** | **Std. Error** | **df** | **t value** | **Pr(>\|t\|)** |
| Intercept | 0.023 | 0.067 | 25 | 0.341 | 0.736 |
| **Linear effect of bin** | **-0.022** | **0.007** | **25** | **-2.946** | **0.007** |
| Quadratic effect of bin | -0.005 | 0.003 | 25 | -1.789 | 0.086 |

**5.2. Fixed effects in comprehension.**

| **LMM for speech comprehension: coherence effect ~ linear bin + quadratic bin + (1\|subj)** | | | | | |
| --- | --- | --- | --- | --- | --- |
|  | **Estimate** | **Std. Error** | **df** | **t value** | **Pr(>\|t\|)** |
| Intercept | -0.013 | 0.101 | 25 | -0.133 | 0.895 |
| Linear effect of bin | -0.002 | 0.011 | 25 | -0.220 | 0.828 |
| **Quadratic effect of bin** | **-0.006** | **0.003** | **25** | **-2.116** | **0.045** |
